# A behavioral paradigm for cortical control of a robotic actuator by freely moving rats in a one-dimensional two-target reaching task

**DOI:** 10.1101/2021.01.24.427325

**Authors:** Syed Muhammad Talha Zaidi, Samet Kocatürk, Tunçer Baykaş, Mehmet Kocatürk

**Affiliations:** Department of Biomedical Engineering, Istanbul Medipol University, Istanbul, Turkey; Department of Computer Engineering, Istanbul Medipol University, Istanbul, Turkey; Health Sciences and Technology Research Institute, Istanbul Medipol University, Istanbul, Turkey; Center for Molecular and Behavioral Neuroscience, Behavioral and Neural Science Graduate Program, Rutgers University-Newark, Newark, NJ, USA; Department of Electrical and Electronics Engineering, Kadir Has University, Istanbul, Turkey

**Keywords:** Motor cortex, neuroprosthetics, brain-machine interface

## Abstract

Controlling the trajectory of a neuroprosthetic device to reach multiple targets is a commonly used brain-machine interface (BMI) task in primates and has not been available for rodents yet. Here, we describe a novel experimental paradigm which enables this task for rats in one-dimensional space for reaching two distant targets depending on their limited cognitive and visual capabilities compared to primates. An online transform was used to convert the activity of a pair of primary motor cortex (M1) units into two robotic actions. The rats were shaped to adapt to the transform and direct the robotic actuator toward the selected target by modulating the activity of the M1 neurons. All three rats involved in the study were capable of achieving randomly selected targets with at least 78% accuracy. A total of 9 out of 16 pairs of units examined were eligible for exceeding this success criterion. Two out of three rats were capable of reversal learning, where the mapping between the activity of the unit pairs and the robotic actions were reversed. The present work is the first demonstration of trajectory-based control of a neuroprosthetic device by rodents to reach two distant targets using visual feedback. The paradigm introduced here may be used as a cost-effective platform for elucidating the information processing principles in the neural circuits related to neuroprosthetic control and for studying the performance of novel BMI technologies using freely moving rats.

## 1. Introduction

Brain-machine interfaces (BMIs) hold great potential for restoration of the motor functions lost due to spinal cord injury or neurodegenerative disorders. Advances in neural interface technology have recently enabled seamless and scar-free tissue integration for the brain-implantable recording probes (Liu et al., 2015; Luan et al., 2017; Zhou et al., 2017; Zhao et al., 2019). Neural decoders allowed monkeys and humans to manipulate multi-degree of freedom robotic arms or cursors on computer screens with their neuronal activity (Velliste et al., 2008; Hochberg et al., 2012; Wodlinger et al., 2015; Flint et al., 2016; Lebedev and Nicolelis, 2017; Nuyujukian et al., 2018). In light of these promising demonstrations, providing a dexterous motor neuroprosthetic control similar to natural movements has been and continues to be a highly active research topic. Mostly monkeys and humans are used to study the performance of neural decoders and the adaptation of the brain to the neuroprosthetic devices (Ganguly et al., 2011; Simeral et al., 2011; Ajiboye et al., 2017; Prins et al., 2017; Li et al., 2020). Even though the cognitive and visual abilities of rodents are limited compared to monkeys and humans (Reinagel, 2015; Zoccolan, 2015), they are also commonly used in BMI research due to the ethical and cost-related considerations. Furthermore, availability of versatile genetic tools for rodents makes them advantageous for studying the neural mechanisms involved in brain adaptation and skill learning (Koralek et al., 2013; Athalye et al., 2018).

The capability of rats to modulate the activity of cortical neurons to obtain rewards has been demonstrated decades ago (Olds, 1965; Hiatt, 1972). Unidirectional manipulation of a brain-controlled robotic actuator has been demonstrated in rats (Chapin et al., 1999) thanks to the advances in microelectrode array technology that enabled multichannel recordings (Nicolelis et al., 1997). Utilization of reinforcement learning-based decoding algorithms allowed the rats to control a robotic arm in three-dimensional space to reach one of two targets (DiGiovanna et al., 2009). Controlling the frequency of an auditory cursor has also been presented using rats for achieving one of two targets (Koralek et al., 2012). The capability of rats to control a robotic actuator unidirectionally and bidirectionally using cortical activity has been demonstrated as well (Arduin et al., 2013, 2014). In these studies, the rats modulated the neural activity to bring water rewards toward themselves using a one-dimensional robotic actuator.

Reaching multiple targets is a typical paradigm applied in BMI experiments with human and monkey subjects for investigating the accuracy of the trajectories generated by the BMI decoders and for studying the neural mechanisms of motor and neuroprosthetic control (Taylor et al., 2002; Carmena et al., 2003; Jarosiewicz et al., 2015). In the task, the subject’s aim is to move a cursor (on a computer screen) or a robotic arm toward multiple distant targets in two or three-dimensional spaces using the visual feedback provided. Implementing this task for rodents in a simpler form depending on their limited cognitive and visual abilities can still provide high throughput and reduce the cost of some BMI studies requiring basic neuroprosthetic actions. On the other hand, the same limitations in the abilities of rats present significant challenges in applying this task even for reaching targets in one-dimensional space such that it has not been demonstrated yet. In this article, we introduce a novel behavioral paradigm which addresses these challenges and enables rats to perform one-dimensional reaching for two distant targets using a cortically controlled robotic actuator. We describe the shaping procedures and the experimental setup we punctiliously designed for rats to this end and investigate the applicability of the present neuroprosthetic control paradigm.

## 2. Methods

### 2.1. Microelectrode Array Implantation

All animal procedures presented in this paper were approved by and conducted in accordance with the regulations of the Istanbul Medipol University Ethics Committee on Animal Maintenance and Experimentation. Three male Wistar rats weighing 400-550 g were chronically implanted bilaterally with two microelectrode arrays in the primary motor cortex (see coordinates below). The arrays were built in-house and each consisted of 16 tungsten microwires (35 μm diameter, polyimide-coated diameter 48 μm; California Fine Wire, CA). The configuration of the microelectrode arrays was 2×8 with 250 μm row and 500 μm column spacing (see **Figure S1**). An additional bare tungsten wire (50 μm diameter) was aligned in parallel with the microwires of the array with 500 μm spacing and used as an indifferent reference electrode. The rats were anesthetized with i.p. injection (1.5 cc/kg) of a mixture of ketamine and xylazine anesthesia (100 mg/kg and 12 mg/kg in saline, respectively). Maintenance of anesthesia was achieved with isoflurane gas as needed. Dexamethasone (0.5 mg/kg i.p.) was administered 1 hour prior to the surgery to attenuate inflammatory response during insertion of the microelectrode arrays (Zhong and Bellamkonda, 2007; Gaire et al., 2018). Craniotomy was created bilaterally and the dura mater was removed over the implantation sites carefully while avoiding damaging blood vessels and the pia mater (Oliveira and Dimitrov, 2008). The pia mater was attached to skull using a 2-octyl and n-butyl cyanoacrylate tissue adhesive (Leukosan, BSN Medical GmbH, Germany) in order to prevent dimpling during lowering the microelectrode arrays (Kralik et al., 2001). The arrays were centered stereotaxically to target forelimb area (AP: +1.5 mm, ML: ±2.5 mm) in both hemispheres (Gioanni and Lamarche, 1985; Kleim et al., 1998) and advanced independently and slowly (50 μm/min) to a cortical depth of ~1200 μm using a hydraulic micropositioner (Narishige MO-82, Japan). When the target depth was achieved, the craniotomy was sealed with a thin layer of cyanoacrylate tissue adhesive and the microelectrode array was fixed to the skull using dental acrylic.

### 2.2. Neural Signal Acquisition

Acquisition and online processing of the neural signals and the control of the components of experimental setup were performed using in-house built Bioinspired Neuroprosthetic Design Environment (BNDE) (Kocaturk et al., 2015). Neural signals were amplified and filtered using 32-channel electrophysiology hardware (PBX Preamplifier, Plexon Inc., TX, USA). The high-cut and low-cut frequencies of the bandpass filter of the hardware were 150 Hz and 8 KHz, respectively, and the passband gain was 1000. Neural signals were acquired with a sampling rate of 31.25 kHz per channel, digitally band-pass filtered (cut-off frequency = 400 Hz–8 kHz) and up-sampled to 62.5 kHz by cubic interpolation. Detection of the neural spikes was performed by applying two amplitude thresholds and spikes were sorted using template matching algorithm based on Bayesian clustering and classification (Lewicki, 1998; Alpaydin, 2010; Kocaturk et al., 2015). A 32-channel commutator (Plexon Inc., TX, USA) was used to keep the headstage cables untangled during electrophysiological recordings.

### 2.3. Rat Operant Conditioning

The experimental setup for cortical control of the robotic actuator is illustrated in **Figure 1** and **Figure S2**. Prior to the initiation of the trials for cortical control of the robotic actuator, the rats were first shaped in the present setup for attending to the robotic workspace. Shaping process was started after a recovery period of four weeks following the microelectrode array implantation surgery. The shaping procedures and the experimental setup introduced previously (DiGiovanna et al., 2009) were employed in this work with some modifications we included for enabling trajectory-based cortical control of the robotic actuator. In this section, we describe the experimental setup and the shaping procedures applied to make the rat attend to the robotic workspace prior to the commencement of cortical control trials.

**Figure 1.**
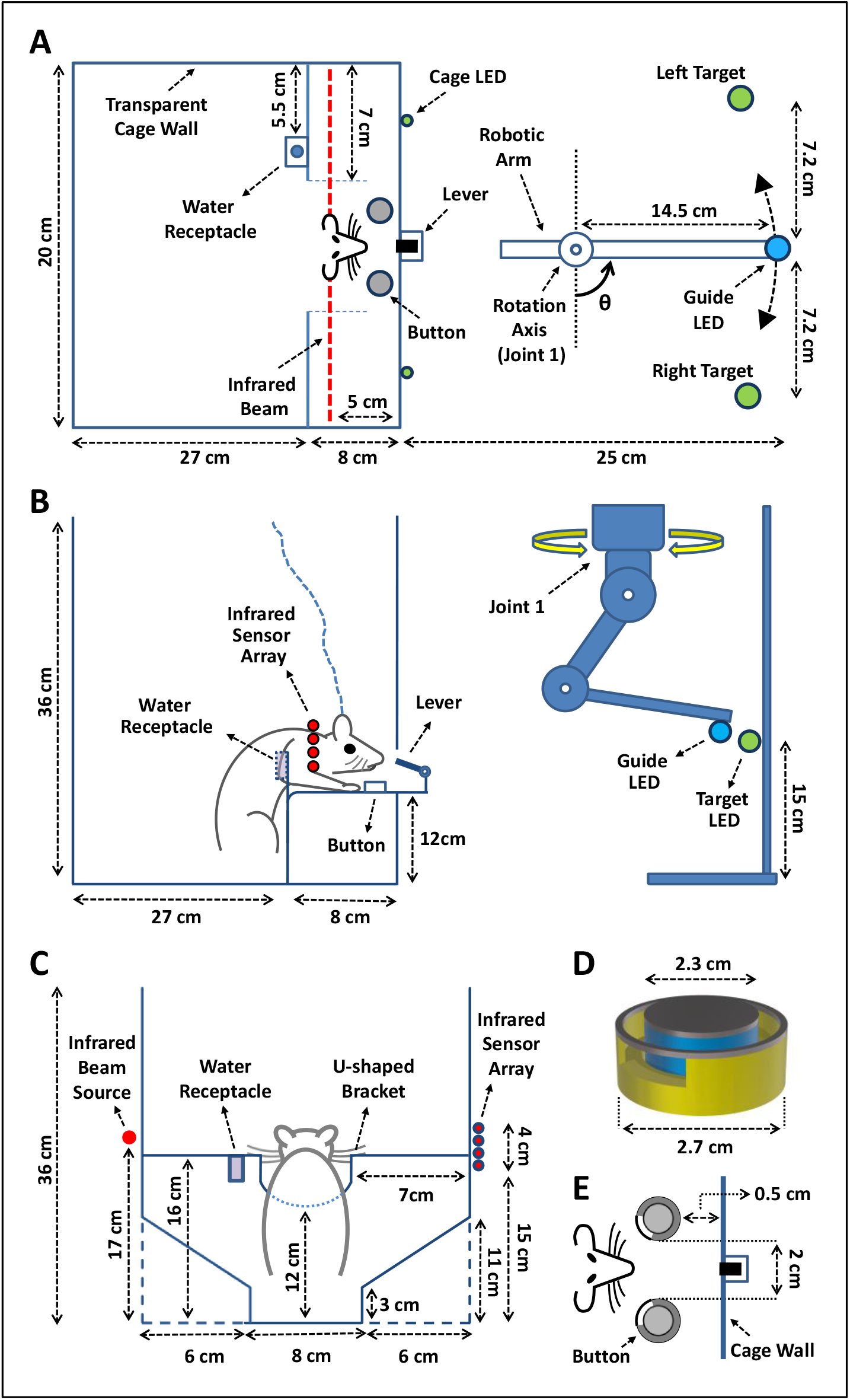
Experimental setup. **(A)** Top view. **(B)** Side view. (**C**) Back view. (**D**) The design of the button which the rats operated with their teeth. The button is surrounded by a cylindrical barrier and an aperture placed onto the side of the barrier such that the rat can insert its lower teeth through the aperture. The rat can press the button with the upper teeth as it puts the lower teeth into the aperture. (**E**) The positions of the buttons and the lever which is used to initiate the trials. The lever is located in front of the rat as shown in (**A**) and (**B**). The orientation of the aperture of each button is adjusted to ease pressing with teeth. During cortical control of the robotic arm, the buttons were covered with a separate lid to disengage the rat from them when needed.

The experimental setup mainly consisted of a behavioral cage and a robotic workspace. The walls of the behavioral cage were transparent so that the robotic workspace was visible to the rat while it was enclosed in the cage **(Figure 1A**). The rats were trained in this environment with a two-button choice task to associate robot control with button pressing. As shown in **Figure 1A,B**, the cage included one lever and two buttons on either side, which were located on a flat platform. The height and the depth of the platform were adjusted according to the size of an adult rat so that it can reach the lever and the buttons while standing on the floor of the cage and putting its forepaws onto the platform as shown in **Figure 1B**. The design of the buttons allowed the rats to operate them by biting with their teeth (see **Figure 1D, E**). Two green cage LEDs were placed onto the front wall of the cage on either side of the platform. An infrared (IR) beam passed through the proximal portion of the platform from the perspective of the rat and reached an array of IR sensors placed onto the opposite wall of the cage. The IR beam was positioned to detect whether the rat’s upper body was over the platform; the trials ended whenever the rat left the platform. A narrow corridor with a width of 8 cm and a height of 3 cm was created on the floor of the cage (see **Figure 1C**) so that the rat always stood at the center of the cage while being capable of reaching the lever and the buttons. The floor of the cage was flat so that the rat could stand comfortably during the trials. A U-shaped bracket (see **Figure 1C**) was placed onto the platform to minimize the variations in posture of the rat during trials and a water receptacle dispensing 15 μl of water was positioned on the wall of the bracket. The amount of water reward was controlled using a solenoid valve. The water delivery by valve generated a click sound during opening and closure, constituting a reward cue for the rat following conditioning. The rats had ad libitum access to food both in the behavioral cage and in their home cage and they were motivated using a 21 hour water deprivation protocol in which they received water only in the behavioral cage during behavioral tasks.

The robotic workspace included a customized version of Lynxmotion AL5D robotic arm (Swanton, VT, USA) and two opposite left/right targets, pointed by green LEDs. The targets were located on the same plane to be reached by the robotic arm moving around its first joint (base servomotor) in one-dimensional space as shown in **Figure 1A,B**. A blue guide LED was mounted onto the tip of the robotic arm as an indicator of the position of the robot endpoint. The elevations of the target LEDs and the guide LED were equal and they were adjusted according to the rat’s field of vision while standing as illustrated in **Figure 1B**. The experimental environment was poorly illuminated to direct rat’s attention toward the LED cues in the robotic workspace. The brightness of the LED cues was adjustable so as to maximize the accuracy in the trials.

Rat training consisted of two phases: The first phase was applied to associate robot control with button pressing and the second phase was used to ensure the rat perceives the position of the robot endpoint **(Figure 2A, B**). In the first training phase, each trial was initiated by the rat by nose-poking the lever in the cage through a hole **(Figure 1B**). With the initiation of a trial, one of the targets was randomly selected and the cage and the target LEDs corresponding to that side were turned on without delay. The guide LED was also illuminated at the onset of each trial. Forty milliseconds after trial initiation, the robot started moving toward the selected target with a constant speed and reached the target within 1.2 s. If the rat pressed the button on the target side 0.7 s after initiation of the robot motion then the trial was considered successful and a water reward was delivered through the receptacle in the cage. The trial was ended without reward delivery, if 1) the rat pressed the wrong button at any time, 2) the rat did not give any response within 1.8 s after robot motion initiation, and 3) the rat pressed the button on the target side earlier than 0.7 s after robot motion initiation. The third condition was applied to prevent the rat from rushing to press the button just after trial initiation and from not attending to the movements of the robot endpoint. Whenever a trial was ended, all LEDs in the experimental setup were turned off and the robotic arm moved back to its default position at the midpoint of the targets to be used in the next trial. At the end of each trial, a refractory period of 2 s was applied to allow the robotic arm to reach its default position and the rat to consume the water reward. The cage LEDs were not lit up in the following trials as the accuracy improved so that the attention of the rat was shifted to the target and guide LEDs in the robotic workspace. When the rat’s accuracy reached the operant conditioning inclusion criterion (i.e. 80% through at least 40 consecutive trials), the first training phase was successfully completed and the second training phase was started. The training paradigm for the second phase is depicted in **Figure 2B**. The entire paradigm was same as the one applied in the first phase except for turning on the target LEDs. In this phase of the training, the only cue for the rat for button choice was the movement direction of Guide LED which constituted the marker of the robot endpoint. After rats reached inclusion criteria of 80% accuracy in this phase of training, the task for cortical control of the robotic actuator commenced.

**Figure 2.**
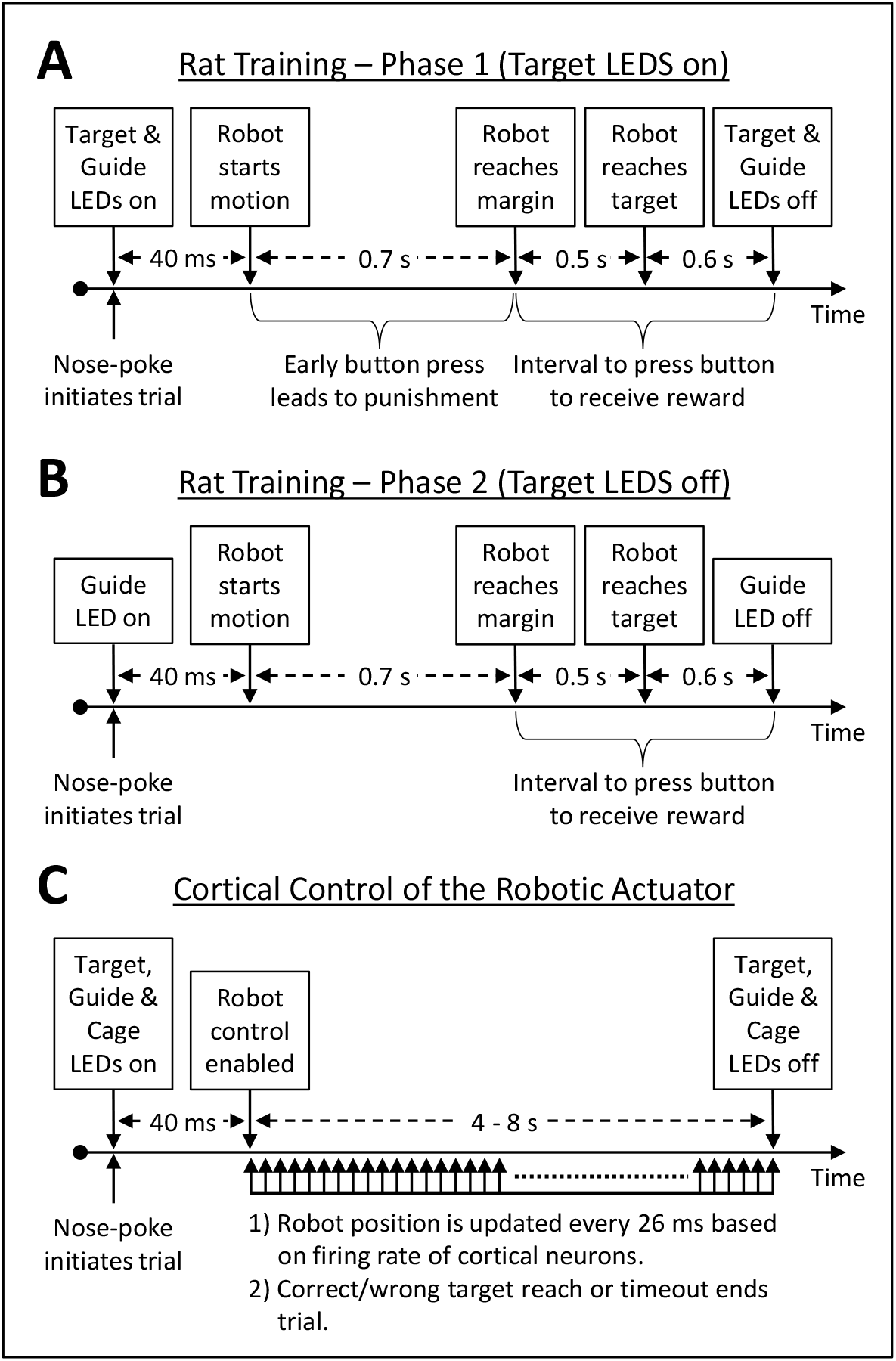
Behavioral paradigm for shaping the rat for neuroprosthetic control. **A**) Paradigm for associating button press with the selected target. (**B**) Paradigm for associating button press with the movement direction of robot endpoint. (**C**) Paradigm for cortical control of the robotic actuator. During cortical control, the transform algorithm converts the firing rates of M1 units into robotic actions.

### 2.4. Cortical Control of the Robotic Actuator

After the rats had been shaped to engage in the robot workspace and associate movements of the robotic actuator with reward (see **Rat Operant Conditioning**), they again entered into the same experimental setup for learning manipulation of the robotic actuator using M1 neurons. In this phase of learning, the movements of the robotic actuator were controlled using the activity of two M1 units rather than being automatic (see **Figure 2C** and **Figure 3A**). A transform algorithm was employed for converting the firing rates of M1 neurons into robotic actions (see **Equation (1)** and **Figure 3A**). The firing rates of the units were calculated by binning the spikes every 26 ms with a sliding 208 ms time window and fed into the below online transform algorithm:

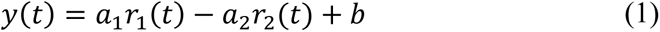

where *r*_1_(*t*) and *r*_2_(*t*) are the firing rates of first and second units at time bin *t. a*_1_ and *a*_2_ are the coefficients and *b* is the bias term used in the transform algorithm where *a*_1_ > 0, *a*_2_ > 0, 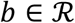. angular velocity of the robotic actuator (*ω*(*t*)) is obtained from the output of the transform (*y*(*t*)) as follows:

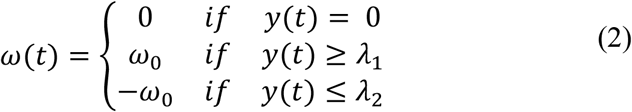

where *λ*_1_ and *λ*_2_ are the firing rate thresholds for movement of the robotic arm toward left or right, respectively, with the constraints *λ*_1_ > 0 and *λ*_2_ < 0. From the perspective of the rat, the robot endpoint moved toward left for the positive values of angular velocity and moved toward right for the negative values **(Figure 1A** and **Figure 3B**). The value of *ω*_0_ was kept constant at 36.76°/s throughout the trials. Therefore, available prosthetic actions were: 1) move left, 2) move right and 3) stay stationary. The angular velocity of the robotic actuator (*ω*(*t*)) was updated every 26 ms, which was also the period for binning the spikes for calculating the firing rates *r*_1_(*t*) and *r*_2_(*t*). An increase in the firing rate of the first unit (*r*_1_(*t*)) directed robotic actuator toward left and an increase in firing rate of the second unit (*r*_2_(*t*)) moved it toward right. In other words, the units opposed each other. The coefficients and the bias term in the transform algorithm (*a*_1_, *a*_2_ and *b*) were manually updated by the experimenter through the trials to guide the rat to acquire both targets by modulating the activity of the units. The absolute values of *λ*_1_ and *λ*_2_ were increased to slow down the rotation of the robotic actuator if the firing rates of the neurons (*r*_1_(*t*) and *r*_2_ (*t*)) were so high that rat’s performance could not be improved through the trials due to rapid movements of the robotic arm.

**Figure 3.**
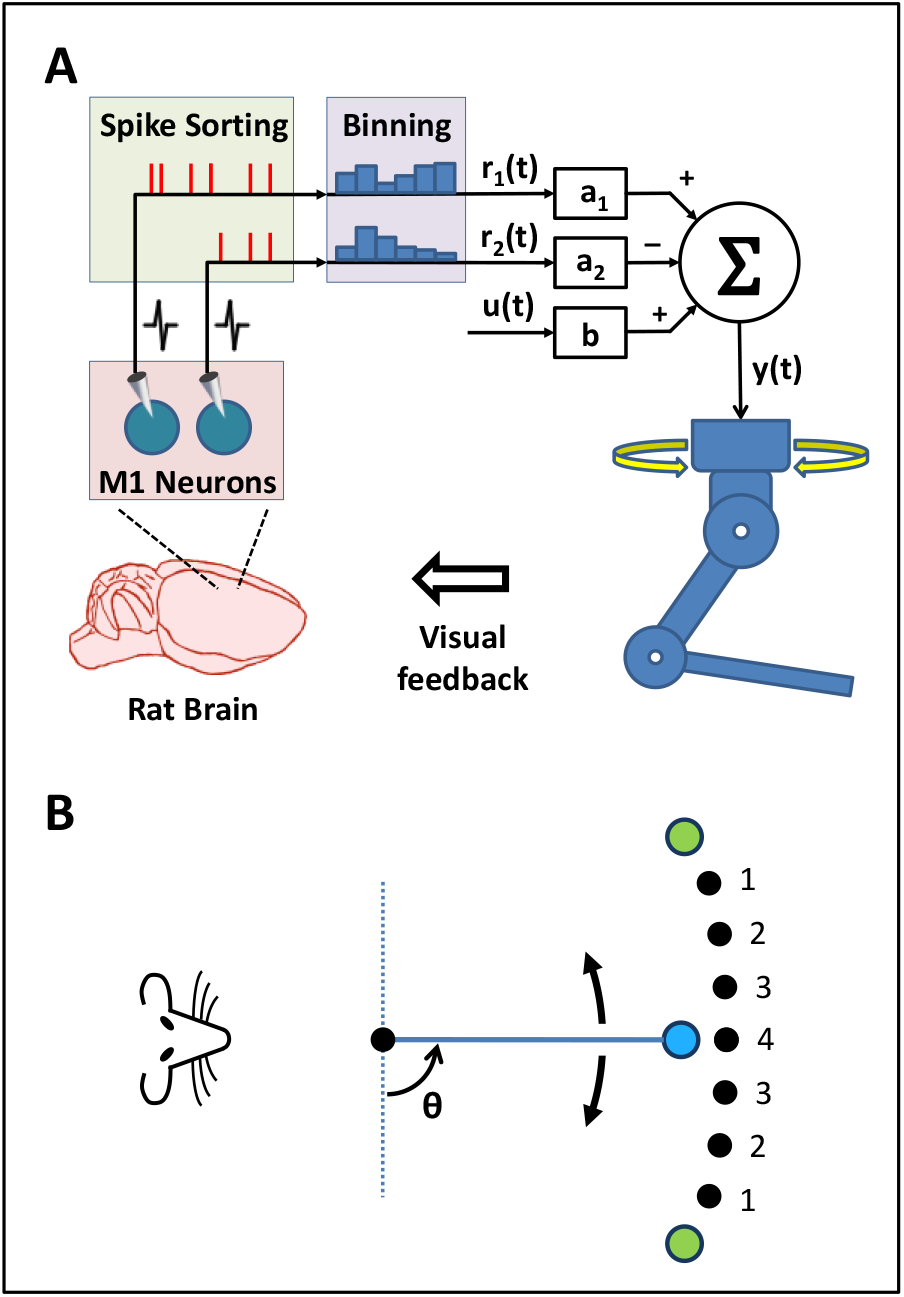
Cortical control of the robotic arm in one-dimensional space. (**A**) The online transform and the neuroprosthetic control architecture. (**B**) The movement of the robot endpoint in one dimensional space. Green circles indicate left/right targets and blue circle denotes the robot endpoint. Black circles represent the initial position of the robot endpoint at the beginning of trials depending on the task difficulty level in effect. The numbers denote the task difficulty levels proportional to the distance between the selected target and the robot endpoint at the beginning of the trials. Level 4 is the highest task difficulty level where the initial position of the robot endpoint is at the midpoint of the targets.

In the present cortical control task, the trials were initiated by the rat by nose-poking as in previous training phases **(Figure 2)**. Forty milliseconds after the trial initiation, the robotic actuator was enabled to move and *r*_1_(*t*) and *r*_2_(*t*) were reset to zero. This delay after trial initiation was introduced to allow the rat to perceive the selected target side and the neural activity to reach baseline after nose-poking. The aim of the rat in the cortical control task was to direct the robotic arm toward the target LED which was randomly selected and illuminated with initiation of the trial. The cage LED on the selected side was also turned on when the trial was started and remained on throughout the trial in order to aid the rat discriminate the target side. Whenever the endpoint of the robotic arm reached the selected target, the rat was rewarded and the trial was classified as “correct”. The trial was ended without reward delivery and classified as “incorrect” if 1) the robotic arm reached the wrong target, 2) none of the targets was achieved within the maximum allowed trial duration (i.e. 4-8 s), 3) all four infrared sensors on the wall of the cage were illuminated by the infrared beam, indicating that the rat left the platform and no longer attended to the robotic workspace. The experimenter was also allowed to manually and immediately terminate the trial without reward delivery by software whenever it was obvious that the target LED was out of the vision of the rat due to the rat’s movements after initiating the trial. Whenever a trial ended, 1) all LEDs in the experimental setup were turned off, 2) the target side in the next trial was determined and 3) the robotic actuator moved to the initial position in the next trial determined according to the task difficulty level in effect and selected target side in that trial (see **Figure 3B**). As in previous training phases, a refractory period of 2 s was applied to allow the robotic actuator to reach the initial position in the next trial and the rat to consume the water reward.

In the cortical control trials, four levels of task difficulty were applied in order to keep the rats motivated in the task and engaged in the robotic workspace. The level of the difficulty determined the distance of the robot endpoint to the selected target at the beginning of a trial. **Figure 3B** illustrates the applied difficulty levels from 1 through 4 and the initial positions of the robot endpoint corresponding to each difficulty level. The task difficulty level was incremented whenever the rat exceeded the cortical control inclusion criterion (i.e. 75% accuracy for 40 consecutive trials in which target side is selected randomly). If the rat could not meet the inclusion criterion through approximately 200 trials, the difficulty level of the task was decremented and the rat was again trained. In the trials in which the robotic arm had varying trajectories, the rat was also manually rewarded by the experimenter when the robotic arm started moving toward the correct target following a movement toward the wrong target. Rewarding the rat whenever it directed the robotic arm toward correct target likely facilitated association of cortical activity with the correct movements of the robotic arm (Thorndike, 1911; Fetz, 1969; Moritz et al., 2008; Arduin et al., 2013). Whenever the rats’ accuracy exceeded inclusion criterion for the highest (4^th^) difficulty level, where the initial position of the robotic actuator is midpoint **(Figure 3B**), the unit pair used was classified as “successful” for neuroprosthetic control. The inclusion criterion in the highest difficulty level was selected as “greater than 75% accuracy for 40 consecutive randomly selected targets”. The probability of exceeding this inclusion criterion by chance is less than 0.001. After reaching the inclusion criterion for the highest task difficulty level, the rats continued using the same units in the rest of the trials in that day until they gave up initiating new trials to receive water. In the following day, a reversal learning task was applied using the same M1 units to determine whether the rats actually adapted to the transform for reaching the selected target or the units involved were stimulated or inhibited directly by the selected target without any learning. To examine this, the sign of the coefficients *a*_1_ and *a_2_* in the transform algorithm were inverted (see **Equation (3)**) so that the mapping between the activity of the units and robotic actions were reversed. In this phase of cortical control task, the rats needed to reverse the activity modulation of the units to direct the robotic actuator toward the correct target; an increase in the firing rate of the first unit (*r*_1_(*t*)) led to a robotic movement toward right rather than left and an increase in the firing rate of the second unit (*r*_2_(*t*)) directed the robotic movement toward left rather than right.

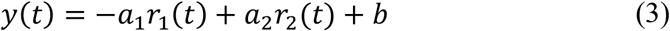

The behavioral paradigm was the same in the reversal learning phase **(Figure 2C)**. The same rules to increase the task difficulty level were applied in this phase of learning. As the rats’ accuracy increased through the trials, the task difficulty level was incremented (see **Figure 3B**). The rat was expected to adapt to the new transform to acquire the selected targets through increasing levels of task difficulty over multiple days of training. When the rat met the inclusion criterion (i.e. >75% accuracy for 40 consecutive trials) for the reversal learning phase in the highest task difficulty level, these units were classified as eligible for reversal learning.

## 3. Results

The rats entered into the experimental setup illustrated in **Figure 1** and they were first trained in the setup for shifting their attention from inside the cage to the robotic workspace. All three rats reached the inclusion criteria in the training phases depicted in **Figure 2A** and **Figure 2B**. The time it took the rats to meet the inclusion criteria to be eligible for starting trials for cortical control of the robotic actuator varied between rats: Rat1 was trained for 16, Rat2 was trained for 22 and Rat3 was trained for 20 days. While 15 μl of water reward at the end of successful trials was sufficient for keeping Rat1 and Rat2 motivated during the tasks, Rat3 required twice the amount of water the other rats needed.

After meeting the operant conditioning inclusion criteria, the rats received the cortical control task (see Figure 2C). The activity and the spike waveforms of the units were reviewed by the experimenter over multiple days prior to commencement of the cortical control task. The cortical control task was started with the first (lowest) difficulty level. Whenever the rats retracted their body through the infrared beam to reach the water receptacle before acquiring the selected target, the trial was immediately terminated without reward delivery. The trials were also ended manually by the experimenter through the software when it was obvious that the rat was not attending to the robotic workspace. In order to help the rat acquire the selected targets, the coefficients in the transform algorithm (**Equation (1)**) were applied differently for each target at the beginning of the training in each difficulty level. For instance, the value of *a*_1_ was increased and the values of *a*_2_ and *b* were decreased when the left target was selected. Inversely, the value of *a*_1_ was decreased and the values of *a*_2_ and *b* were increased when the right target was selected. As the rats’ accuracy improved, the difference between the values of these coefficients applied differently for each target was reduced gradually. Finally, the same coefficients and bias terms were applied for both targets. When the inclusion criterion was exceeded for both targets with the finalized values of *a*_1_, *a*_2_ and *b*, the difficulty level was incremented and the optimal values for the parameters of the transform were searched again in the next task difficulty level by repeating the same shaping approach. Selection of the same target for several consecutive trials during the training sessions also facilitated reinforcing the neural activity modulations leading to acquisition of the selected targets. Randomization in target selection was introduced afterwards as the target acquisition performance improved through a series of trials in which same target was selected consecutively.

The trajectory of the robotic actuator in a trial with highest (4^th^) difficulty level is shown in Figure 4. In this representative trial, the right target was randomly selected and the rat acquired the selected target by modulating the activity of the pair of the units involved in the neuroprosthetic control. The initial position of the robot endpoint in this difficulty was at the midpoint of the targets (see **Figure 3B**). The coefficients in the transform were *a*_1_ = 1, *a*_2_ = 1 and *b* = 4.8 Hz, the firing rate thresholds were *λ*_1_ =4.8 Hz and *λ*_2_ =4.8 Hz during the control of the robotic actuator for reaching both left and right targets in this task difficulty level. By the initiation of a trial via nose poke, the robotic actuator was enabled to move and the firing rate values of the units (r(*r*_1_(*t*) and *r*_2_(*t*)) were reset to zero **(Figure 4C,D**). The output of the transform (*y*(*t*)) was calculated throughout the trial **(Figure 4E**). In **Figure 4A,B**, we can see the activity of Unit 1 was mostly higher than that of Unit 2 just before and at the beginning of the trial and the robot endpoint moved toward left by the trial initiation. As the right target LED was selected for illumination in this trial, the rat modified the firing rates of the units and directed the robotic actuator toward right **(Figure 4F)**. One can see the inertia of the robotic arm from the difference between the delivered pulse width commands (green colored trace) and actual trajectory of the robotic arm (orange colored trace) **(Figure 4F)**. The selected target (right target) was acquired within 2.23 s based on the actual trajectory of the robotic actuator and the rat was rewarded with water at the end of trial.

**Figure 4.**
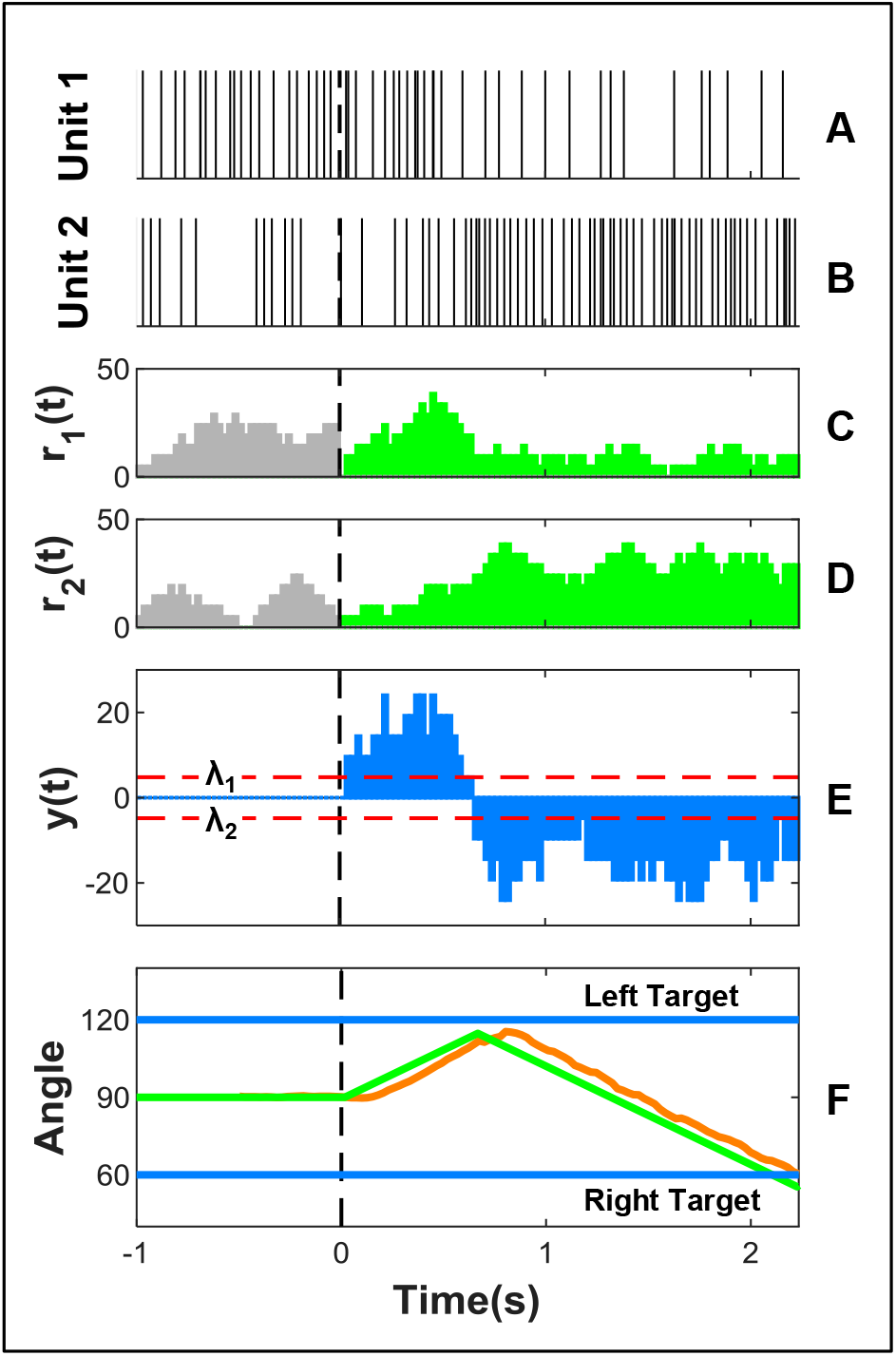
Change in the joint angle of the robotic actuator based on the firing rates of the unit pair involved in neuroprosthetic control. (**A** and **B**) The raster of the spikes generated by the units involved in control. (**C** and **D**) The firing rates of the units (***r*_1_**(***t***) and ***r_2_***(***t***)), which are fed to the transform (**Equation (1)**). Firing rates of the units before/after the initiation of the trial are shown by gray/green bars, respectively. (**E**) The output of the transform (***y***(***t***)) used in the manipulation of the robotic actuator. Horizontal dashed red lines indicate the values of thresholds ***λ*_1_** and ***λ*_2_** used in **Equation (2)**. (**F**) The trajectory of the robotic actuator. The green trace indicates the joint angle values sent to the base servomotor of the robotic arm and the orange trace indicates its actual trajectory. Horizontal blue lines present the positions of the left and right targets in terms of the joint angle corresponding to the base servomotor of the robotic arm. Vertical dashed black lines in all plots represent the time on which the robot control was enabled by a nose poke.

The overall target reach accuracy of Rat1 through 50 consecutive trials is illustrated in **Figure 5A.** The highest (4^th^) task difficulty level was applied through these trials. The target reach accuracy of the rat through these trials was 90%. The rat was not rewarded in 5 out of 50 trials since the correct target was not achieved. In one of these trials (18^th^ trial), the robotic actuator reached the wrong target. In remaining four, the rat moved toward the water receptacle and left the platform earlier than acquisition of any target. This action of the rat was detected using the infrared beam located in the cage **(Figure 1A,B)** and the trial was ended immediately. The control of the robotic actuator through these 50 trials is shown in **Movie S1**. The resulting target reach accuracy was achieved within 6 days of training by gradually incrementing the task difficulty level (see **Figure 3B**). The number of trials performed per day ranged from 521 to 1175. As depicted in **Figure 5A,** the finalized values of the parameters of the transform in the highest task difficulty level were *a*_1_=1, *a*_2_=1 and *b*=4.8 Hz. The firing rate thresholds used in the transform algorithm were *λ*_1_=4.8 Hz and *λ*_2_ =4.8 Hz. The changes in the joint angle of the robotic arm for the correct trials are illustrated in **Figure 5C**. The trajectory through 8^th^ trial in **Figure 5A** is also shown in **Figure 4. Figure 6A** demonstrates the raster plot of the spikes generated by the unit pair involved in cortical control of the robotic actuator. After the target acquisition performance had reached plateau levels in the 4^th^ (highest) task difficulty level, the ongoing session on that day was continued until the rat gives up initiating new trials.

**Figure 5.**
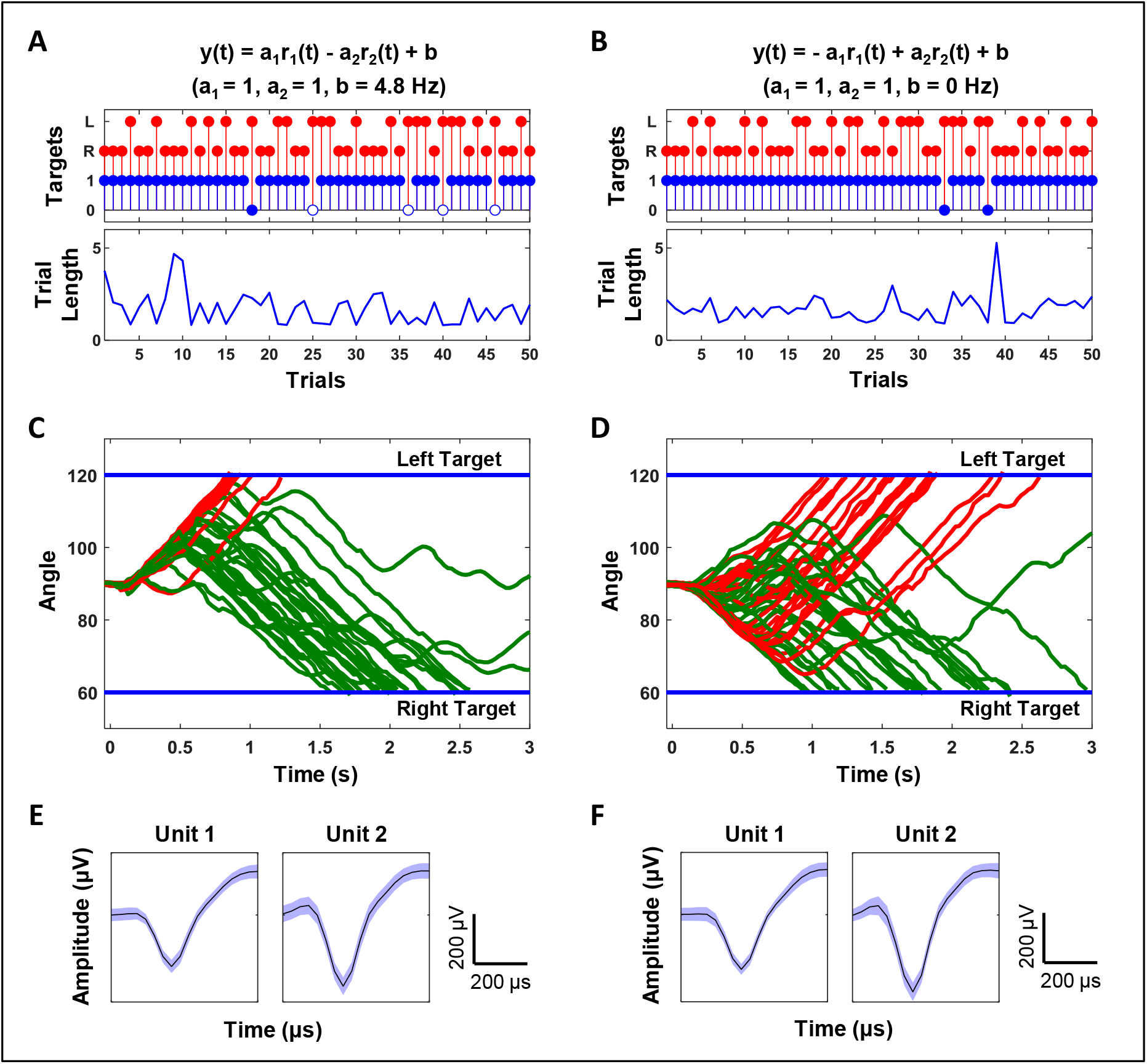
Target reach performance of Rat1 using the same M1 units before and after introducing reversal learning. (**A**) Target reach accuracy and the lengths of trials when the transform algorithm presented with **Equation (1)** is applied. The selected target for each trial is represented by red stems (L for left trial and R for right trial) and the blue stems show if the correct (selected) target was acquired (1) or not (0). Blue stems with empty markers indicate the trial was ended without reward delivery since the rat left the platform before target reach and consecutively the infrared beam sensor array in the cage (see **Figure 1A,B**) was activated. Blue stems with filled markers for the unsuccessful trials point the trial was ended due to the acquisition of the wrong (opposite) target. (**B**) Target reach accuracy and the lengths of trials after reversal learning with application of the transform algorithm given with **Equation (3)**. The bottom graphs in (**A**) and (**B**) present the lengths of the trials in terms of seconds. (**C** and **D**) The trajectories of the robotic arm through the correct trials depicted in (**A**) and (**B**), respectively. Red/green traces denote the trajectories for the selected left/right targets in terms of the joint angle corresponding to the base servomotor of the robotic arm. (**E** and **F**) Average of waveforms of 100 spikes for M1 units used in the trials shown in (**A**) and (**B**), respectively. The shaded regions present the standard deviation.

**Figure 6.**
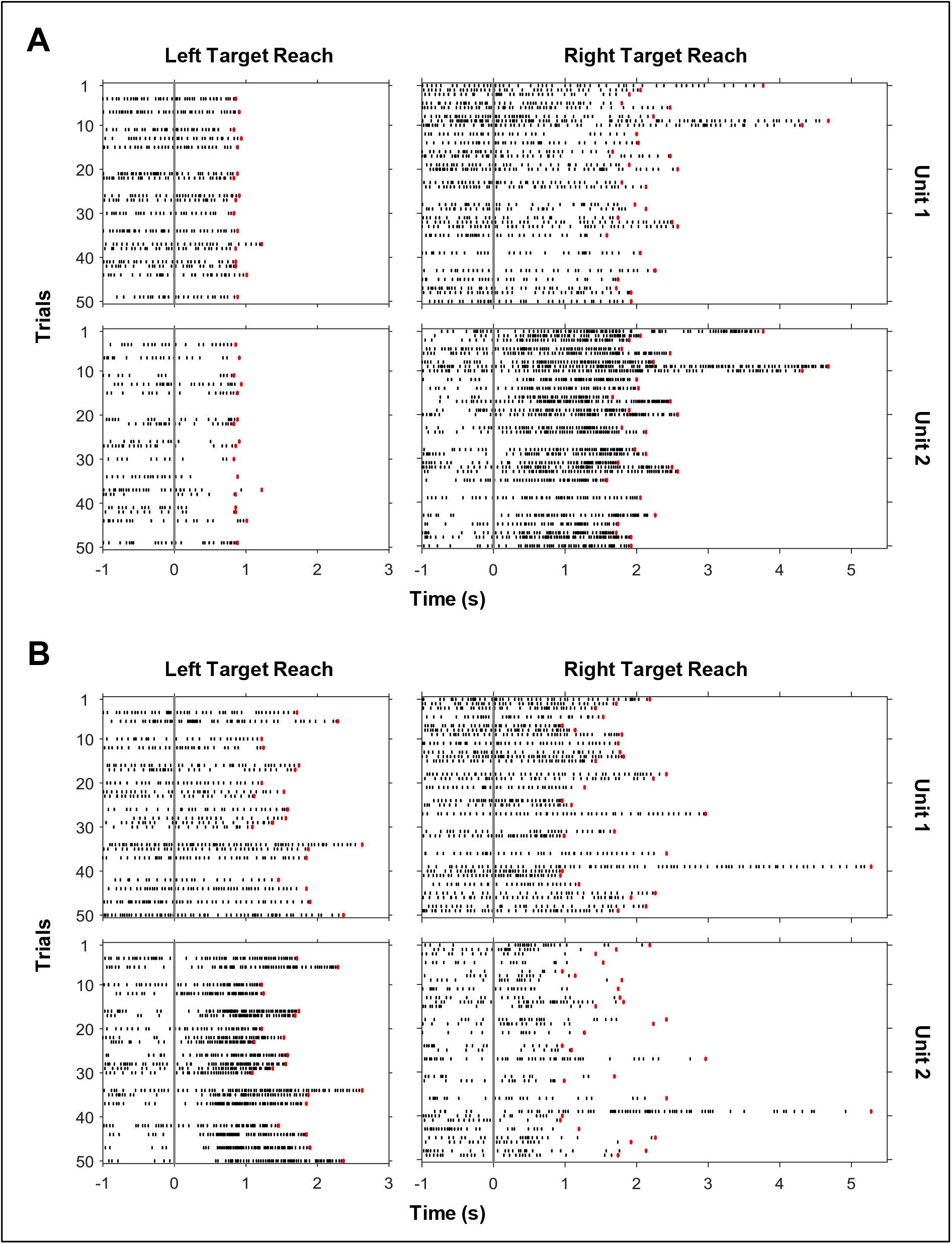
Activity pattern of the units involved in the trials presented in Figure 5. (**A**) Raster plot for the trials depicted in **Figure 5A,C.** (**B**) Raster plot for the trials depicted in **Figure 5B,D.** Only the trials ended with the achievement of the correct target are presented in the graphs. Red markers indicate the target achievement times. Vertical gray lines denote the time on which the robot control was enabled. Each row of graphs corresponds to the unit shown on the rightmost side of the row. Each column of graphs corresponds to the selected target side.

After reaching the inclusion criterion for the highest difficulty level (i.e. greater than 75% correct target achievement through at least 40 consecutive trials in which the targets are selected randomly), the reversal learning task was initiated in the following day using the same M1 units. In the reversal learning task, the signs of the coefficients of the transform algorithm were inverted (**Equation (3)**) so that the rat was to adapt to the new transform to achieve the randomly selected targets again. **Figure 5B** illustrates the target achievement performance of Rat1 for 50 consecutive trials in the reversal learning task after reaching plateau levels in accuracy. **Figure 6B** presents raster plot of the activity of the unit pair involved in neuroprosthetic control through these trials. The target reach accuracy of the rat was 95%. The rat was not rewarded in 2 out of 50 trials since it acquired the wrong targets. **Movie S2** demonstrates the manipulation of the robotic arm by Rat1 through these 50 trials. The present target achievement performance was obtained within 2 days of training by gradually incrementing the task difficulty level. The traces in **Figure 5D** present the changes in the joint angle of the robotic arm through the correct trials in the reversal learning task. As presented in **Figure 5B**, **t**he finalized values of the parameters of the transform in the highest task difficulty level in the reversal learning task were *a*_1_=1, *a*_2_=1 and *b*=0 Hz. The firing rate thresholds applied in the transform algorithm were *λ*_1_ =4.8 Hz and *λ*_2_ =4.8 Hz.

A total of 16 pairs of units from three rats were examined for the neuroprosthetic control. **Table 1** provides a brief summary of the target reach performance of the rats using these unit pairs. The criteria for selection of the units for cortical control task were the waveform stability over multiple days prior to commencement of cortical control task, having clearly identifiable waveforms in spike sorting and having a firing rate more than 1 Hz while the forepaws of the rat are placed onto the platform shown in **Figure 1B**. Based on these criteria the number of selected single units varied between rats. Rat1 had 5 units, Rat2 had 9 units and Rat3 had 10 units. 4 pairs of units were created for Rat1, 5 pairs were created for Rat2 and 7 pairs were created for Rat3 for cortical control of the robotic arm. The hemispheres and numbers of the microelectrode array channels through which the units were isolated are listed for each pair in **Table 1** (see column “Pairs” in Table 1). For example, a pair of units was formed using the units isolated through 4^th^ channel of the microelectrode array implanted into the left hemisphere (L4) and 10^th^ channel of the microelectrode array implanted into the right hemisphere (R10) in Rat3 for the first cortical control session. If the task difficulty level could not be increased for a pair over multiple days of training, one or both of the units in the pair were replaced with other units isolated through the neural recordings. In Table 1, the pairs of units with which the rats were not capable of reaching the inclusion criterion are indicated by cross marks (**×**). The pairs with which the rat met the inclusion criterion are demonstrated with the percentage of reaching correct targets through 40 consecutive trials (see column “Acc.” in Table 1). The percentages of acquiring the correct targets in the reversal learning task are also listed in the second row allocated for the relevant unit pair. The pairs with which the inclusion criterion was not achieved in the reversal learning task are marked with a dash (–). For instance, the accuracy for pair R11-R15 in Rat1 in reversal learning task was 95% and the pair R6 -R12 in same rat was not eligible for reversal learning as indicated by a dash (–). The numbers of days of training in the cortical control mode required to reach the listed accuracies are also presented in the table (see column “Days” in Table 1). The values of the transform algorithm parameters (*a*_1_, *a*_2_, *b*, *λ*_1_ and *λ*_2_) enabling reaching the listed accuracies are given in the table in the relevant columns. Briefly, a total of 9 out of 16 pairs allowed reaching the inclusion criterion for acquiring the randomly selected targets in the highest (4^th^) task difficulty level. 4 out of 9 successful pairs (i.e. R11-R15, R10- R12, L14–L15 and L4-L6) additionally allowed reaching the inclusion criterion in the reversal learning task in the highest task difficulty level. The trajectory, target reach accuracy and raster of spikes for the pair R11-R15 in Rat1 are illustrated in **Figure 5** and Figure 6. 7 out of 16 pairs (pointed by cross (**×**) marks) were ineligible for presenting activity modulations for effective control of the prosthetic actuator. For the pairs L4-R7 and R4-R12 in Rat2, the stability of the waveforms for spike sorting diminished during the cortical control training sessions. For the pair L5-L7 in Rat2, the reliability of the spike sorting was reduced during the reversal learning task and further training for this pair was ended. Only unit pair R4-L1 in Rat 2 remained well-isolated during the reversal learning task and this pair appeared to be ineligible for reversal learning in the present control paradigm.

**Table 1.**
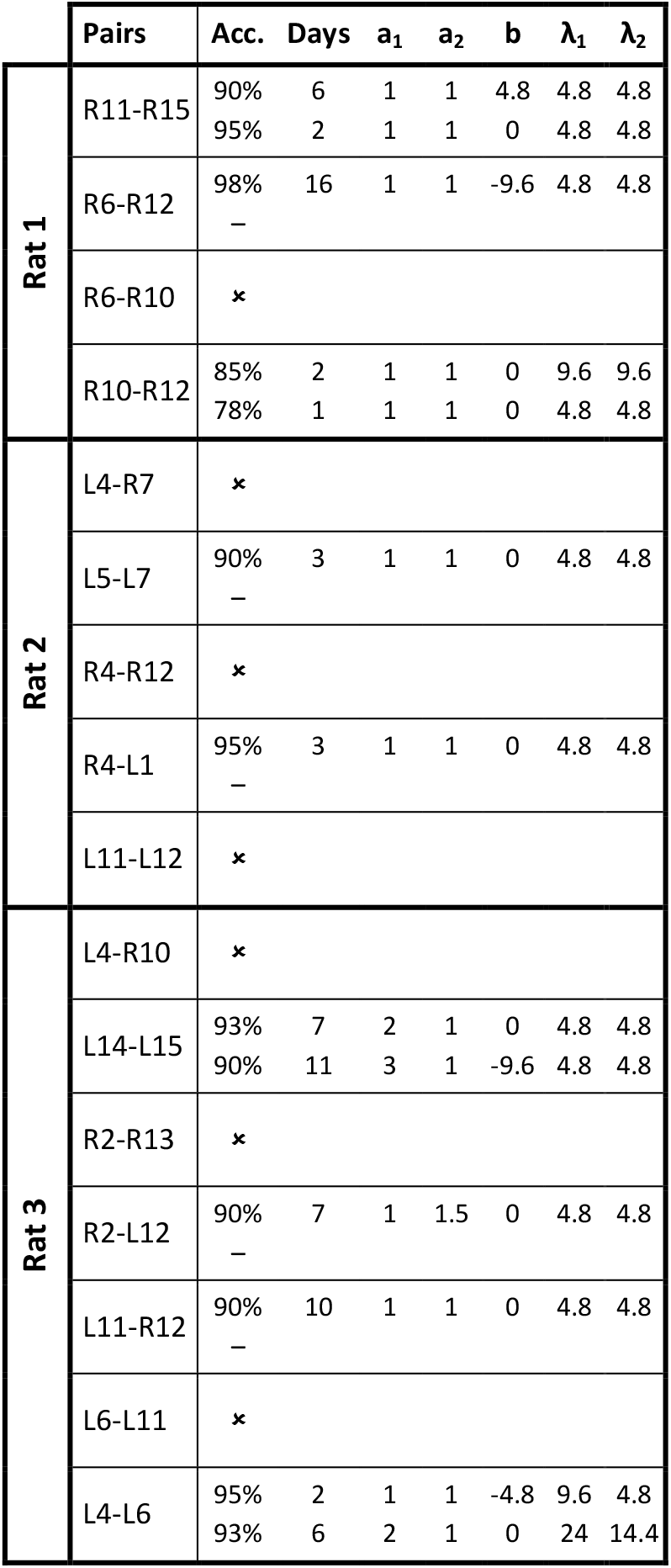
Target reach performance using M1 units.

## 4. Discussion

In the present work, our goal was to develop a behavioral paradigm and an experimental setup that allow the use of rats for motor neuroprosthetic control in a two-target reaching task. The solution described here enabled all three rats involved in the study to control a robotic arm through M1 neurons for reaching two opposite targets in one-dimensional space. In the present cortical control task, an online transform algorithm was employed to make two M1 units oppose each other and convert their firing rates into two robotic actions (see **Equation (1)**). The rats learned modulating the firing rates of the units and selecting the robotic actions directing the robot endpoint toward the indicated targets. The adaptation of the rats to the transform algorithm was also validated by reversing the mapping between the unit pairs and the robotic actions (see **Equation (3)**).

The components and the structure of the experimental setup were specifically designed for enabling freely moving rats to control a motor neuroprosthesis in a one-dimensional two-target reaching task. Importantly, the dimensions of the platform onto which the rat put its forepaws during trials (see **Figure 1B**), the dimensions of the U-shaped bracket on the edge of platform (see **Figure 1C**) and the width of the corridor on the floor of the cage (see **Figure 1C**) were adjusted according to the size of the rat in order to minimize the variations in the posture through neuroprosthetic control trials. These measures were crucial in neuroprosthetic control since the activity patterns of the recorded neurons and visual feedback from the robotic workspace might vary from trial to trial due to the discrepancies in the posture and such variations in the posture might negatively affect the learning curve of the rat. Secondly, in the design of the behavioral cage, the water receptacle which provided the rewards was placed onto the wall of the U-shaped bracket (see **Figure 1A,C**) rather than somewhere close to buttons and lever in the cage. We chose this location to eliminate the probability of the rats being distracted by the water receptacle during neuroprosthetic control trials. Thirdly, the narrow U-shaped bracket placed onto the edge of the platform **(Figure 1C**) allowed the rats to reach the water receptacle in a stereotypical manner at the end of the neuroprosthetic control trials. This design allowed us to set a precise criterion for ending the trials without reward delivery whenever the rat left the platform prior to target acquisition. This movement of the rat (leaving the platform) could be practically detected using the infrared beam shown in **Figure 1A-C** and the trials could be automatically and immediately ended without releasing reward (see **Figure 5A**). Thus, the rats could be enforced to keep their upper body in the U-shaped bracket and above the platform throughout the online control of the robotic arm. In addition, we used LEDs with adjustable brightness to indicate the position of the robot endpoint and the selected target, and optimized their brightness to maximize the rats’ visual abilities and accuracies through the trials. Consequently, the overall design of the behavioral setup guided the rats to receive visual feedback from the robotic workspace during neuroprosthetic control.

Other key components in the design of the cage were involved in the training process prior to the cortical control trials; two buttons located on either side of the platform (see **Figure 1D,E**) were used to move the attention of the rats to the robotic workspace through the training phases depicted in **Figure 2A,B**. The buttons were specifically designed to allow the rats to operate them by doing timely, stereotypical and intentional motions in response to environmental cues (Skinner, 1938; Dickinson, 1994; Jin and Costa, 2010; Kawai et al., 2015). These buttons have been suitable to be used by the rats over the course of the shaping process prior to the initiation of the cortical control task. In the present experimental setup, the time required to shape the rats for attending to the robotic workspace prior to cortical control task ranged from 16 to 22 days.

After associating the button presses with the cues in the robotic workspace, the rats entered into the cortical control task illustrated in **Figure 2C** and **Figure 3A**. As listed in Table 1, the time required to exceed the success criterion in cortical control task ranged from 2 to 16 days. The values of the coefficients and the bias term used in the transform (**Equation (1)**) varied based on the baseline firing rates and modulation characteristics of the units involved in neuroprosthetic control. The values of the parameters in the transform were gradually updated by the experimenter by trial-and-error based on the target achievement performance of the rats through the cortical control trials. Simultaneously, the rats adapted with the transform and improved their target reach accuracy by trial-and-error, in other words by reinforcement learning. The task difficulty level was increased gradually over the course of several days of training and the rats were shaped toward the most difficult task. Finally, the rats reached the success criterion in the highest task difficulty level and the values of the parameters in the transform converged. In the present work, the process of updating the parameters of the transform was manually performed by the experimenter. Automatizing this process can be a matter of developing novel, adaptive BMI decoders where the parameters are updated according to the target reach accuracy of the subject (Mahmoudi et al., 2013; Pohlmeyer et al., 2014; Kocaturk et al., 2015). In addition, we believe the performance of BMI decoders in general can also be studied using the present experimental setup after enabling the subjects to control the robotic actuator using two isolated units as described in this work.

The paradigm presented here allowed the rats to move freely during the cortical control of the robotic arm. During neuroprosthetic control we monitored and recorded the behavior using two infrared cameras without introducing distracting stimuli. The setup allowed us to visually and qualitatively observe the movements of the rat during the trials (see **Movie S1- S2**). It is known that changes in cortical activity during neural operant conditioning can be accompanied by motor or muscle activity (Hiatt, 1972; Fetz and Baker, 1973). The limitation of our study was the lack of usage of the tools (e.g. electromyography (EMG) electrodes implanted into the muscles) for quantitative analysis of variations in the correlations between cortical and muscle activity. Previously it has been demonstrated that the pitch of an auditory cursor can be intentionally controlled using M1 units without any significant change in the EMG activity (Koralek et al., 2012). Based on our qualitative visual observations, the movements of the rat during neuroprosthetic control ranged from no movement to overt body movements varied from trial to trial, from unit pair to unit pair involved in neuroprosthetic control (Fetz and Baker, 1973).

In the present paradigm, the acquisition of the targets by the robotic actuator was the predictor of reward in contrast to the previous demonstrations where the subject brought the reward toward itself using a neuroprosthetic actuator (Chapin et al., 1999; Arduin et al., 2013, 2014). The rats in our study became capable of controlling the movements of the reward predictor (the robot endpoint) rather than reward itself (e.g. water, sucrose solution or food). In addition, the rat was directly involved in the control of the trajectory of the robot endpoint throughout a robotic reaching task in contrast to the goal-based BMIs where the BMI controller learned the necessary trajectory to reach the selected target in three-dimensional space (DiGiovanna et al., 2009; Mahmoudi and Sanchez, 2011). Lastly, in the present control paradigm the rats manipulated the position of a robot’s endpoint using visual feedback instead of shifting the frequency of a tone using auditory feedback, which had been an effectively employed neuroprosthetic control paradigm in previous studies to reach one of two target tones (Koralek et al., 2012, 2013). In this context, the rat behavioral paradigm presented here is novel and similar to the trajectory-based, center-out cortical control of a cursor commonly achieved by monkey or human subjects in BMI studies. The control of the movement of a sipper tube in two dimensions via manipulating a joystick has also been demonstrated in rats previously (Slutzky et al., 2010). In light of this study future work may assess the capability of rats to control the trajectory of a neuroprosthetic device in two dimensions. In addition, practicality of a cursor on a PC monitor as a neuroprosthetic device (Simeral et al., 2011) can also be studied in rats provided that the brightness and contrast levels obtained in our work using LEDs can also be achieved by the monitor to be used.

## 5. Conclusion

We described a behavioral paradigm along with an experimental setup meticulously designed for shaping freely moving rats to acquire two distant targets using a brain-machine interface (BMI). A BMI task similar to the one applied with primates was demonstrated in a simpler form for rats despite the difficulties caused by the limitations in their cognitive abilities. In contrast to the existing techniques applied in rats for studying BMI control, the present setup enabled them to perform trajectory-based, center-out control of a robotic actuator in one-dimensional space using the visual feedback provided. In the control paradigm, the firing rates of a pair of cortical units were fed into an online transform algorithm whose output was used as a control signal for manipulation of the robotic actuator. The task difficulty level represented the distance between the initial position of the robot endpoint and the selected target. By gradually increasing the task difficulty level and manually updating the parameters of the transform through the training sessions, the rats were effectively shaped to control the robotic actuator to acquire the randomly selected targets starting from a central position. We believe the experimental setup introduced here can provide a cost-effective and practical alternative for studying the practicality of novel BMI technologies and open opportunities for advancing our understanding of the neural basis of neuroprosthetic control.

## Supplementary Material

**Movie S1.** Cortical control of the robotic arm during 50 consecutive trials listed in **Figure 5A** (back view).

**Movie S2.** Cortical control of the robotic arm during 50 consecutive trials listed in **Figure 5A** (front view).

**Movie S3.** Cortical control of the robotic arm during 50 consecutive trials listed in **Figure 5B** (back view).

**Movie S3.** Cortical control of the robotic arm during 50 consecutive trials listed in **Figure 5B** (front view).

## Acknowledgment

This research was supported by The Scientific and Technological Research Council of Turkey (TÜBİTAK), Grant #115E257 and #117E286.

## Notes

### Competing Interest Statement

The authors have declared no competing interest.

## References

Ajiboye, A. B., Willett, F. R., Young, D. R., Memberg, W. D., Murphy, B. A., Miller, J. P., et al. (2017). Restoration of reaching and grasping movements through brain-controlled muscle stimulation in a person with tetraplegia: a proof-of-concept demonstration. Lancet 389, 1821–1830. doi:10.1016/S0140-6736(17)30601-3.

Alpaydin, E. (2010). Introduction to machine learning. 2nd ed. Cambridge, Mass.: MIT Press.

Arduin, P. J., Frégnac, Y., Shulz, D. E., and Ego-Stengel, V. (2013). “Master” neurons induced by operant conditioning in rat motor cortex during a brain-machine interface task. J. Neurosci. 33, 8308–20. doi:10.1523/JNEUROSCI.2744-12.2013.

Arduin, P. J., Frégnac, Y., Shulz, D. E., and Ego-Stengel, V. (2014). Bidirectional control of a one-dimensional robotic actuator by operant conditioning of a single unit in rat motor cortex. Front. Neurosci. 8, 206. doi:10.3389/fnins.2014.00206.

Athalye, V. R., Santos, F. J., Carmena, J. M., and Costa, R. M. (2018). Evidence for a Neural Law of Effect. 1029, 1024–1029. doi:10.1126/SCIENCE.AAO6058.

Carmena, J. M., Lebedev, M. A., Crist, R. E., O’Doherty, J. E., Santucci, D. M., Dimitrov, D. F., et al. (2003). Learning to Control a Brain–Machine Interface for Reaching and Grasping by Primates. PLoS Biol. 1, E42. Available at: http://www.ncbi.nlm.nih.gov/pubmed/14624244.

Chapin, J. K., Moxon, K. a, Markowitz, R. S., and Nicolelis, M. A. L. (1999). Real-time control of a robot arm using simultaneously recorded neurons in the motor cortex. Nat. Neurosci. 2, 664–70. doi:10.1038/10223.

Dickinson, A. (1994). “Instrumental Conditioning,” in Animal Learning and Cognition, ed. N. J. Mackintosh (Orlando: Academic Press), 45–79.

DiGiovanna, J., Mahmoudi, B., Fortes, J., Principe, J. C., and Sanchez, J. C. (2009). Coadaptive brain-machine interface via reinforcement learning. IEEE Trans. Biomed. Eng. 56, 54–64. doi:10.1109/TBME.2008.926699.

Fetz, E. E. (1969). Operant conditioning of cortical unit activity. Science 163, 955–8. Available at: http://www.ncbi.nlm.nih.gov/pubmed/4974291 [Accessed November 20, 2014].

Fetz, E. E., and Baker, M. A. (1973). Operantly conditioned patterns on precentral unit activity and correlated responses in adjacent cells and contralateral muscles. J. Neurophysiol. 36, 179–204. doi:10.1152/jn.1973.36.2.179.

Flint, R. D., Scheid, M. R., Wright, Z. A., Solla, S. A., and Slutzky, M. W. (2016). Long-Term Stability of Motor Cortical Activity : Implications for Brain Machine Interfaces and Optimal Feedback Control. 36, 3623–3632. doi:10.1523/JNEUROSCI.2339-15.2016.

Gaire, J., Lee, H. C., Hilborn, N., Ward, R., Regan, M., and Otto, K. J. (2018). The role of inflammation on the functionality of intracortical microelectrodes. J. Neural Eng. 15, 066027. doi:10.1088/1741-2552/aae4b6.

Ganguly, K., Dimitrov, D. F., Wallis, J. D., and Carmena, J. M. (2011). Reversible large-scale modification of cortical networks during neuroprosthetic control. Nat. Neurosci. 14, 662–7. doi:10.1038/nn.2797.

Gioanni, Y., and Lamarche, M. (1985). A Reappraisal of Rat Motor Cortex Organization by Intracortical Microstimulation.

Hiatt, D. E. (1972). Investigations of operant conditioning of single unit activity in the rat brain.

Hochberg, L. R., Bacher, D., Jarosiewicz, B., Masse, N. Y., Simeral, J. D., Vogel, J., et al. (2012). Reach and grasp by people with tetraplegia using a neurally controlled robotic arm. Nature 485, 372–5. doi:10.1038/nature11076.

Jarosiewicz, B., Sarma, A. A., Bacher, D., Masse, N. Y., Simeral, J. D., Sorice, B., et al. (2015). Virtual typing by people with tetraplegia using a self-calibrating intracortical brain-computer interface. Sci. Transl. Med. 7, 313ra179–313ra179. doi:10.1126/scitranslmed.aac7328.

Jin, X., and Costa, R. M. (2010). Start/stop signals emerge in nigrostriatal circuits during sequence learning. Nature 466, 457–462. doi:10.1038/nature09263.

Kawai, R., Markman, T., Poddar, R., Ko, R., Fantana, A. L., Dhawale, A. K., et al. (2015). Motor Cortex Is Required for Learning but Not for Executing a Motor Skill. Neuron, 1–13. doi:10.1016/j.neuron.2015.03.024.

Kleim, J. a, Barbay, S., and Nudo, R. J. (1998). Functional reorganization of the rat motor cortex following motor skill learning. J. Neurophysiol. 80, 3321–5. Available at: http://www.ncbi.nlm.nih.gov/pubmed/9862925.

Kocaturk, M., Gulcur, H. O., and Canbeyli, R. (2015). Toward Building Hybrid Biological/in silico Neural Networks for Motor Neuroprosthetic Control. Front. Neurorobot. 9, 8. doi:10.3389/fnbot.2015.00008.

Koralek, A. C., Costa, R. M., and Carmena, J. M. (2013). Temporally precise cell-specific coherence develops in corticostriatal networks during learning. Neuron 79, 865–72. doi:10.1016/j.neuron.2013.06.047.

Koralek, A. C., Jin, X., Long, J. D., Costa, R. M., and Carmena, J. M. (2012). Corticostriatal plasticity is necessary for learning intentional neuroprosthetic skills. Nature 483, 331–5. doi:10.1038/nature10845.

Kralik, J. D., Dimitrov, D. F., Krupa, D. J., Katz, D. B., Cohen, D., and Nicolelis, M. A. L. (2001). Techniques for Chronic, Multisite Neuronal Ensemble Recordings in Behaving Animals. Methods 25, 121–150. doi:10.1006/meth.2001.1231.

Lebedev, M. A., and Nicolelis, M. A. L. (2017). Brain-Machine Interfaces: From Basic Science to Neuroprostheses and Neurorehabilitation. Physiol. Rev. 97, 767–837. doi:10.1152/physrev.00027.2016.

Lewicki, M. S. (1998). A review of methods for spike sorting: the detection and classification of neural action potentials. Netw. Comput. Neural Syst. 9, R53–78. Available at: http://www.ncbi.nlm.nih.gov/pubmed/10221571 [Accessed October 9, 2014].

Li, W., Ji, S., Chen, X., Kuai, B., He, J., Zhang, P., et al. (2020). Multi-source domain adaptation for decoder calibration of intracortical brain-machine interface. J. Neural Eng. 17, 066009. doi:10.1088/1741-2552/abc528.

Liu, J., Fu, T.-M., Cheng, Z., Hong, G., Zhou, T., Jin, L., et al. (2015). Syringe-injectable electronics. Nat. Nanotechnol. doi:10.1038/nnano.2015.115.

Luan, L., Wei, X., Zhao, Z., Siegel, J. J., Potnis, O., Tuppen, C. A., et al. (2017). Ultraflexible nanoelectronic probes form reliable, glial scar–free neural integration. Sci. Adv. 3, e1601966. doi:10.1126/sciadv.1601966.

Mahmoudi, B., Pohlmeyer, E. A., Prins, N. W., Geng, S., and Sanchez, J. C. (2013). Towards autonomous neuroprosthetic control using Hebbian reinforcement learning. J. Neural Eng. 10, 066005. doi:10.1088/1741-2560/10/6/066005.

Mahmoudi, B., and Sanchez, J. C. (2011). A symbiotic brain-machine interface through value-based decision making. PLoS One 6, e14760. doi:10.1371/journal.pone.0014760.

Moritz, C. T., Perlmutter, S. I., and Fetz, E. E. (2008). Direct control of paralysed muscles by cortical neurons. Nature 456, 639–42. doi:10.1038/nature07418.

Nicolelis, M. A. L., Ghazanfar, a a, Faggin, B. M., Votaw, S., and Oliveira, L. M. (1997). Reconstructing the engram: simultaneous, multisite, many single neuron recordings. Neuron 18, 529–37. Available at: http://www.ncbi.nlm.nih.gov/pubmed/9136763.

Nuyujukian, P., Albites Sanabria, J., Saab, J., Pandarinath, C., Jarosiewicz, B., Blabe, C. H., et al. (2018). Cortical control of a tablet computer by people with paralysis. PLoS One 13, e0204566. doi:10.1371/journal.pone.0204566.

Olds, J. (1965). Operant conditioning of single unit responses. Excerpta Medica Int. Congr. Ser. 87, 372–380.

Oliveira, L. M. O., and Dimitrov, D. (2008). “Surgical Techniques for Chronic Implantation of Microwire Arrays in Rodents and Primates,” in Methods for Neural Ensemble Recordings, ed. M. A. L. Nicolelis (Boca Raton: CRC Press).

Pohlmeyer, E. A., Mahmoudi, B., Geng, S., Prins, N. W., and Sanchez, J. C. (2014). Using reinforcement learning to provide stable brain-machine interface control despite neural input reorganization. PLoS One 9, e87253. doi:10.1371/journal.pone.0087253.

Prins, N. W., Pohlmeyer, E. A., Debnath, S., Mylavarapu, R., Geng, S., Sanchez, J. C., et al. (2017). Common marmoset (Callithrix jacchus) as a primate model for behavioral neuroscience studies. J. Neurosci. Methods 284, 35–46. doi:10.1016/j.jneumeth.2017.04.004.

Reinagel, P. (2015). Using rats for vision research. Neuroscience 296, 75–9. doi:10.1016/j.neuroscience.2014.12.025.

Simeral, J. D., Kim, S.-P., Black, M. J., Donoghue, J. P., and Hochberg, L. R. (2011). Neural control of cursor trajectory and click by a human with tetraplegia 1000 days after implant of an intracortical microelectrode array. J. Neural Eng. 8, 025027. doi:10.1088/1741-2560/8/2/025027.

Skinner, B. F. (1938). The behavior of organisms: an experimental analysis. New York: Appleton-Century-Crofts.

Slutzky, M. W., Jordan, L. R., Bauman, M. J., and Miller, L. E. (2010). A new rodent behavioral paradigm for studying forelimb movement. J. Neurosci. Methods 192, 228–32. doi:10.1016/j.jneumeth.2010.07.040.

Taylor, D. M., Tillery, S. I. H., and Schwartz, A. B. (2002). Direct cortical control of 3D neuroprosthetic devices. Science 296, 1829–32. doi:10.1126/science.1070291.

Thorndike, E. L. (1911). Animal intelligence: experimental studies. New York: The Macmillan Company, doi:10.5962/bhl.title.55072.

Velliste, M., Perel, S., and Spalding, M. (2008). Cortical control of a prosthetic arm for self-feeding. Nature 453, 1098–101. doi:10.1038/nature06996.

Wodlinger, B., Downey, J. E., Tyler-Kabara, E. C., Schwartz, A. B., Boninger, M. L., and Collinger, J. L. (2015). Ten-dimensional anthropomorphic arm control in a human brain-machine interface: difficulties, solutions, and limitations. J. Neural Eng. 12, 016011. doi:10.1088/1741-2560/12/1/016011.

Zhao, Z., Li, X., He, F., Wei, X., Lin, S., and Xie, C. (2019). Parallel, minimally-invasive implantation of ultra-flexible neural electrode arrays. J. Neural Eng. 16, 035001. doi:10.1088/1741-2552/ab05b6.

Zhong, Y., and Bellamkonda, R. V. (2007). Dexamethasone-coated neural probes elicit attenuated inflammatory response and neuronal loss compared to uncoated neural probes. Brain Res. 1148, 15–27. doi:10.1016/j.brainres.2007.02.024.

Zhou, T., Hong, G., Fu, T.-M., Yang, X., Schuhmann, T. G., Viveros, R. D., et al. (2017). Syringe-injectable mesh electronics integrate seamlessly with minimal chronic immune response in the brain. Proc. Natl. Acad. Sci. 114, 5894–5899. doi:10.1073/pnas.1705509114.

Zoccolan, D. (2015). Invariant visual object recognition and shape processing in rats. Behav. Brain Res. 285, 10–33. doi:10.1016/j.bbr.2014.12.053.

